# Prognostic Significance of preoperative serum CA125, CA19-9, CA72-4, CEA, and AFP in Patients with Endometrial cancer

**DOI:** 10.1101/2024.01.23.576857

**Authors:** Zi-hao Wang, Yun-zheng Zhang, Shu-wen Ge, Luhe-Shan, Bo Wang, Zi-yu Zhang, Qi-jun Wu, Xiao-xin Ma

## Abstract

**Objective:** To determine preoperative serum CA125, CA19-9, CA72-4, CEA, and AFP with prognostic value, and to establish a risk score based on CA125, CEA, AFP levels for predicting the overall survival (OS) and progression-free survival (PFS) of endometrial cancer (EC) patients.

**Methods:** A retrospective cohort study with 2081 EC patients was conducted at Shengjing Hospital of China Medical University. Patient baseline information, tumor characteristics, and data on five serum biomarkers (CA125, CA19-9, CA72-4, CEA, and AFP) were collected. Hazard ratios (HRs) and 95% confidence intervals (CIs) were determined using univariate or multivariate Cox proportional hazard models. log-rank test and Kaplan-Meier analysis were used to compared survival, Data were randomly divided into a training cohort (50%, N = 1041) and an external validation cohort (50%, n = 1040). the least absolute shrinkage and selection operator (Lasso)-Cox regression model was used to screen the independent factors for establishing risk score. And develop nomograms for survival rate prediction.

**Results:** Multivariate analysis showed Elevated CA125 (P<0.0001) AFP (P <0.0001) and CEA(P=0.037) were identified as independent biomarkers for PFS. Increased CA125 (P = 0.003) AFP (P <0.0001) and CEA(P=0.014) were independent factors associated with OS. CA125, AFP and CEA were thus incorporated in an innovative Risk score (RS) by Lasso-Cox regression model, The RS was also an independent indicator for PFS (P<0.0001) and OS (P<0.0001). Furthermore, we developed and validated nomogram based on Cox regression models. The discriminative ability and calibration of the nomograms revealed good predictive ability, as indicated by the calibration plots.

**Conclusion:** This study suggests that the risk score based on preoperative serum levels of CA125, CEA, and AFP was prognostic biomarkers for predicting progression-free survival and overall survival for EC patients. Nomograms based on the RS and clinicopathological features accurately predict Prognosis of EC patients.

## 1. Introduction

Endometrial cancer (EC) is one of the most common gynecological cancer in developed countries such as Europe and the United States, accounts for approximately 20% to 30% of all gynecological tumors. Recent years has also seen year-on-year increases in tumor incidence in Asian countries(*1, 2*). In 2021, there were 66,570 new cases of EC in the United States and 12,940 deaths related to it(*3*). With increases in life expectancy and rates of obesity, corresponding increases in frequency of EC worldwide are expected (*4*), EC ranks second among the gynecological malignancies in China(*5*). An early symptom of EC is postmenopausal vaginal bleeding, most patients can be diagnosed early. However, there are still more than 28% of patients are diagnosed at an advanced stage(*6*), And 10–20% of early-stage EC patients have recurrence or metastasis after treatment, which has become the main cause of their death(*7*). Currently, potential factors to determine prognosis (tumor grade, staging) are being closely monitored in an effort to predict EC patient survival. However, some patients presenting with similar clinicopathological features experience very different outcomes(*8*). As such, we sought to identify effective EC biomarkers that able to predict prognosis in EC patients and identify potential patients at high risk of developing EC.

A tumor marker refers to substance produced and secreted by tumor cells that subsequently enter body fluids or tissues(*9*). At present, the mechanisms of tumorigenesis and development are not completely clear, and identification of tumor markers is paramount. With recent developments in the field of tumor biology and immunology, some serum tumor markers that may potentially make a significant impact on the initiation and progression of gynecological malignancies have been discovered(*10–12*). High-risk populations and women with systemic and advanced disease may benefit from early detection using serum markers. Sensitive and specific Tumor serum markers would also provide a means of monitoring treatment response and enable prediction of recurrence or metastasis.

Five serological tumor markers, including carbohydrate antigen 125 (CA125), carbohydrate antigen 19-9 (CA19-9), carbohydrate antigen 72-4 (CA72-4), carcinoembryonic antigen (CEA), and alpha-fetoprotein (AFP) are routinely used in clinical practice as tools for cancer diagnosis and prediction of prognosis. The most widely used biomarker for EC is CA125. It is used to early diagnose, detect tumor recurrence, and to predict survival outcomes(*13–16*). However, the ability of CA125 to specifically reflect the metastatic potential of tumors is still insufficient. CA125 levels will not increase during the progression of some patients. Moreover, Numerous EC serological tumor markers have been studied, including HE4 (Human epididymal protein 4), CA19-9, AFP, and CEA(*17, 18*). Serum HE4 can be used as an independent prognostic factor for patients with EC(*19*). Moreover, the predictive efficacy of HE4 for disease recurrence seems to be superior to CA125(*20*), especially in high-risk EC patients with high incidence of distant metastasis(*21*). Bian et al. showed that combined use of HE4, CA125, CA72-4, and CA19-9 is valuable in the early diagnosis of EC and that these markers are useful immunological tissue markers for patients with EC (*22*). Unfortunately, serum markers are more commonly used for evaluation of diagnosis for EC patients. identification of other specific serum cancer biomarkers is important clinically for the early diagnosis of recurrence or metastasis.The study of EC tumor markers and their expression pattern can provide additional information when making judgments about EC pathology and course of treatment. It plays a significant role in patient evaluation and progress of individualized treatment, it is also especially crucial for prediction of prognosis. We performed a retrospective study at Shengjing Hospital of China Medical University. The aim was to assess the relationship between pre-operative serological tumor biomarkers and clinicopathological parameters in EC patients, and determine novel biomarkers that are associated with post-operative survival.

However, the predictive ability of single-serum tumor markers remains insufficient. We aim to explore a new signature which involve multiple serum tumor markers, may improve prognosis prediction

## 2. Materials and Methods

### 2.1 Study population

We performed a retrospective study of EC patients from December 2010 to March 2020 at Shengjing Hospital affiliated to China Medical University. This study was approved by the Institutional Review Committee of the Shengjing Hospital of China Medical University (No. 2017PS292K). All patients included in the study met the following criteria: 1) age≥18 years, 2) underwent hysterectomy at Shengjing Hospital, 3) post-operative diagnosis of EC by pathology, 4) no history of other malignant tumors, 5) no active infection, 6) no blood disease or other serious comorbidities, 7) no thromboembolic events, and 8) complete data for all required variables in this study. A total of 1286 patients did not meet all the inclusion criteria and were excluded, and 2081 patients were included (Fig. 1). Informed consent was acquired from all participants of research.

**Figure 1.** Flow chart of the patient inclusion and exclusion process.

### 2.2 Sample-size calculation

The sample size was calculated using an online sample calculation website, http://riskcalc.org:3838/samplesize/. According to previous research, abnormality of AFP level had the lowest prevalence and had a prevalence of approximately 1.8%. We expected that patients with high AFP levels would have an HR value of 3, compared to those with normal AFP levels. The Type I error rate set in this study was 0.05, the degree of certainty of the study was 0.8, and the approximate ratio of the sample size between the three populations was 1. A minimum sample size of 1680 was calculated for this study.

### 2.3 Data collection

The following demographic and clinical data were collected from the electronic medical records of the Shengjing Hospital Information System: patient’s age at diagnosis, pre/post-menopausal status, International Federation of Obstetrics and Gynecology Association (FIGO) stage, tumor grade, histology type, tumor myometrial invasion depth, and existing comorbidities. Pre-operative levels of serum tumor markers (including CA125, CA72-4, CA19-9, AFP, and CEA) in all enrolled patients was measured using a chemiluminescence immunoassay.

Data collection was conducted by professional gynecologists and pathologists. Evaluation and classification of tumors was in accordance with the WHO classification during post-operative examination of tumor pathology, and all tumors were clinically staged according to FIGO guidelines (2009). Pathologists divide tumors into well (G1), moderately (G2), or poorly (G3) differentiated, and tumor histology was classified as type I (mainly endometrioid) or type II (non-endometrioid).

### 2.4 Follow-up and outcome

Post-operative follow-up was conducted over the telephone. The date of last follow-up was Dec 30, 2022. The primary outcome used for assessment was OS and PFS. The date of death was obtained during follow-up or from a provided death certificate.

### 2.5 Statistical analysis

The data input was checked to ensure its accuracy. All data were statistically analyzed using SPSS ver. 20.0 (IBM Corp, Armonk, NY). Descriptive statistics were used to represent patients’ general characteristics. The chi-square test and the t-test were respectively used for qualitative and quantitative variables to identify associations between clinicopathological features and tumor biomarkers. A Kaplan-Meier analysis was used to compare differences in survival, and a log-rank test was used to make comparisons between sub-groups. Univariate and multivariate analyses were used to identify factors that related to patient survival, and the Cox proportional hazards model was used to test for independence of the effect. Data were expressed as hazard ratios (HRs) and 95% confidence intervals (CIs). Further Lasso regression analysis was performed to obtain a risk score formula based on serum expression levels of the three tumor markers. Based on the clinical prediction model, the prognostic histogram was constructed using R version 4.2.0. The C index is used to assess the predictive power of the model. and calibration curve was used to evaluate the calibration degree of the clinical prediction model.

## 3. Results

### 3.1 Patient Baseline Characteristics

Characteristics of eligible patients are summarized in Table 1. Of the 2081 patients, 1964 had tumors classified as pathological type I, and 117 had tumors classified as type II. 600 patients had not experienced menopause at the time of diagnosis, and the remaining 1481 cases were diagnosed with endometrial cancer after menopause. Post-operative FIGO staging by examination of EC pathology identified 1645 stage I cases, 436 stage II-IV cases. 1052 patients had highly differentiated tumors, 1029 patients had moderately and poorly differentiated tumors. The seroprevalence rates of pre-operative CA125, CA19-9, CA72-4, AFP, and CEA were 27.3% (569/2081), 17.2% (357/2081), 25.7% (534/2081), 1.73% (36/2081), and 5.09% (106/2081) respectively.

**Table 1:**
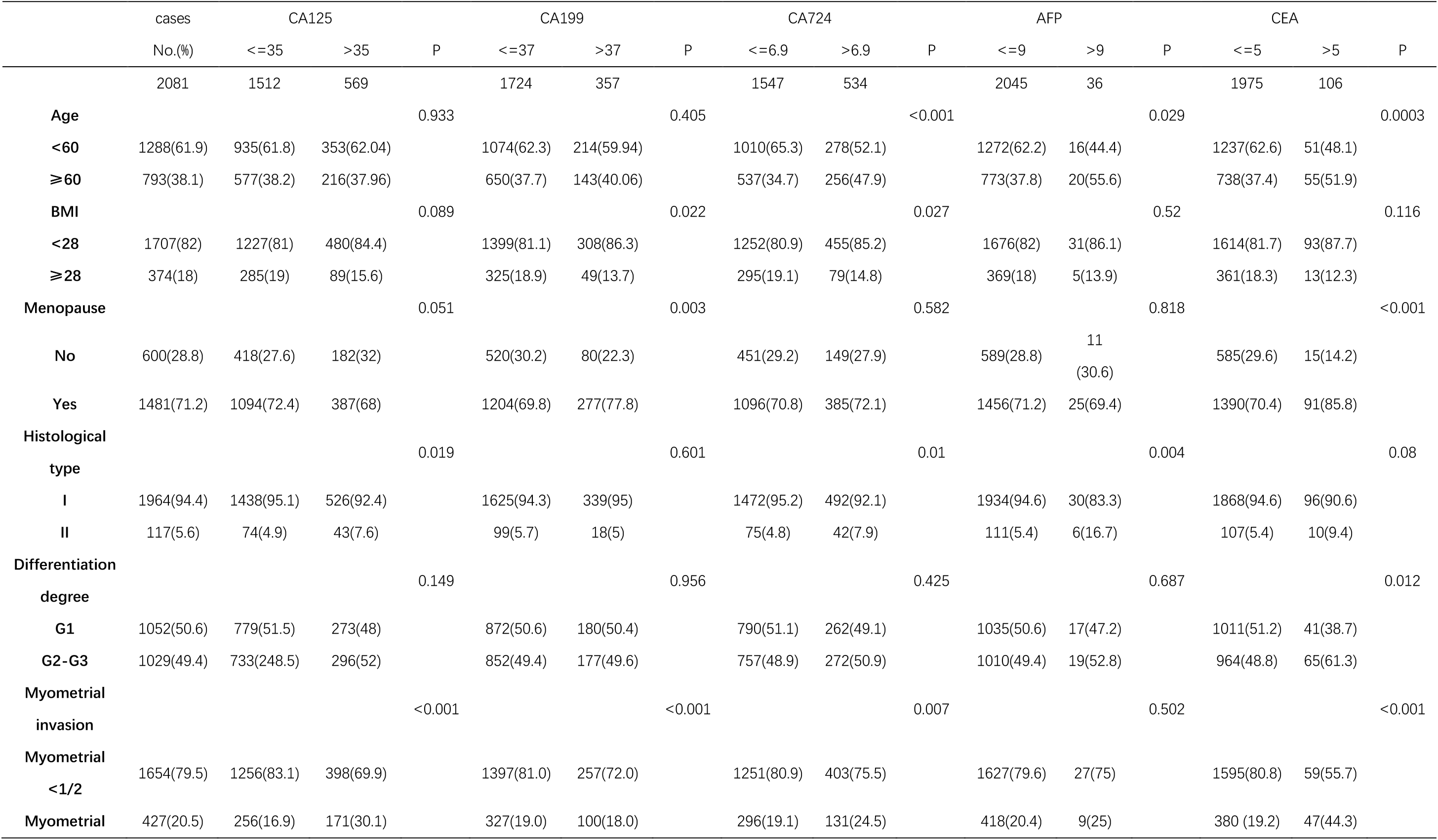
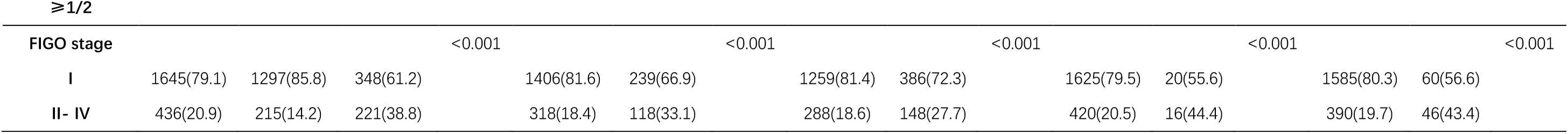
Patient characteristics and relationships between the clinicopathological characteristics and serum CA125, CA19-9, CA72-4, AFP and CEA.

### 3.2 Serum CA125, CA19-9, CA72-4, CEA, AFP levels for patients wth EC

Classified patients into two groups, event (recurrence/metastasis/death occurred) and non-event (none of the above occurred) groups. The two groups were compared for levels of pre-operatively measured CA125, CA19-9, CA72-4, CEA, and AFP. Levels of CA125 (P < 0.0001), CA19-9 (P = 0.0005), CA72-4 (P = 0.0021), CEA (P <0.0001), and AFP (P <0.0001) were significantly higher in the event group compared to the non-event group (Fig. 2). As such, CA125, CA19-9, CEA, and AFP may be used as potential markers to predict the prognosis of EC patients.

**Figure 2.** Comparison of preoperative serum CA125, CA19-9, CA72-4, AFP and CEA levels in event (recurrence/metastasis/death occurred) and non-event groups of endometrial cancer patients (N = 906). ∗ We defined P <0.05 as statistical difference.

### 3.3 Univariate and Multivariate Survival Analysis of EC Patients

For this study, the median follow-up time was 48 months, 5-year OS was 90.0%, and 5-year PFS was 87.0%. Single factor survival analysis was performed to determine the effect of variables such as age, menopausal status, pathological type, degree of differentiation, FIGO stage, depth of myometrial invasion, hypertension, diabetes, complications, lymphovascular invasion (LVSI), expression of ER, PR, P53 in pathological tissues, serum CA125, CA19-9, CEA, and AFP levels in EC patients.

In univariate analysis (Table 2), age≥60 (P < 0.001), menopausal status (P < 0.001), pathological typeⅡ (P < 0.001), low degree of differentiation (P < 0.0001), FIGO stage≥Ⅱ (P < 0.001), depth of myometrial invasion ≥ 1/2(P < 0.001), Positive lymphovascular infiltrates(P < 0.001), Negative ER expression(P < 0.001), Negative PR expression(P < 0.001) and Positive P53 expression(P =0.006), CA125>35 (P < 0.001), CA72-4>6.9 (P = < 0.001), CEA>5 (P < 0.001), and AFP>9 (P =< 0.001) were significantly associated with poor OS prognosis for EC patients. Serum levels of CA125 (P < 0.001), CA72-4 (P = < 0.001), CEA (P < 0.001), and AFP (P =< 0.001) were also associated with poor PFS prognosis for EC patients. Differences in survival for serum CA125, CA19-9, CA72-4, CEA, AFP were compared using a Kaplan-Meier analysis. We identified serum levels of CA125, CA72-4, AFP and CEA as factors for prognosis of OS and PFS in EC patients (Fig. 3)

**Figure 3.** The Kaplan-Meier statistical analysis was used to analyze the overall survival curves and the progression-free survival curves of patients with different levels of each biomarker. (A-i): Kaplan–Meier curves of OS for different groups of CA125 level; (A-ii): Kaplan–Meier curves of PFS for different groups of CA125 level; (B-i): Kaplan–Meier curves of OS for different groups of CA19-9 level; (B-ii): Kaplan–Meier curves of PFS for different groups of CA19-9 level; (C-i): Kaplan–Meier curves of OS for different groups of CA72-4 level; (C-ii): Kaplan–Meier curves of PFS for different groups of CA72-4 level; (D-i): Kaplan–Meier curves of OS for different groups of AFP level; (D-ii): Kaplan–Meier curves of PFS for different groups of AFP level; (E-i): Kaplan–Meier curves of OS for different groups of CEA level; (E-ii): Kaplan–Meier curves of PFS for different groups of CEA level.

**Table 2:**
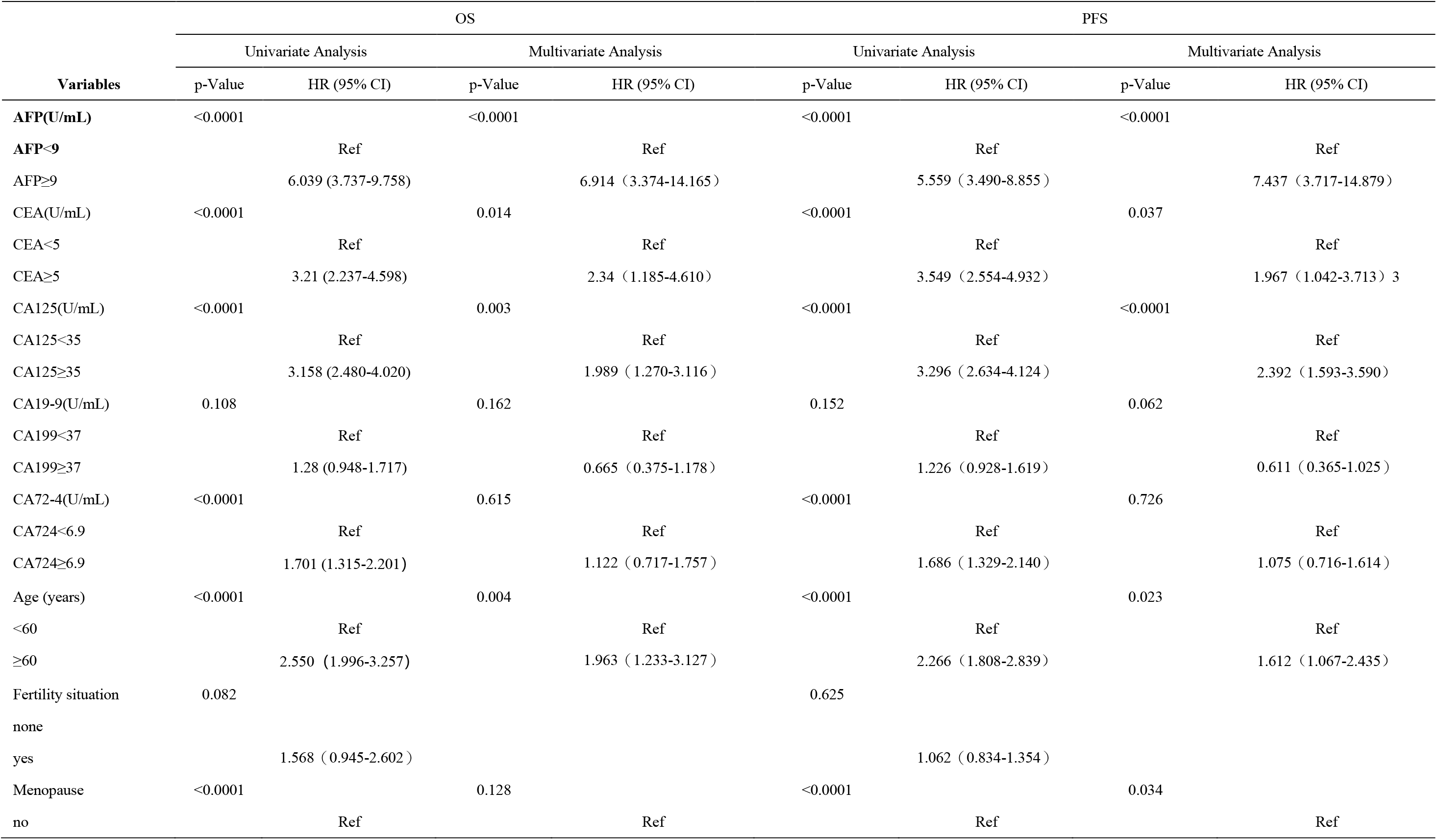
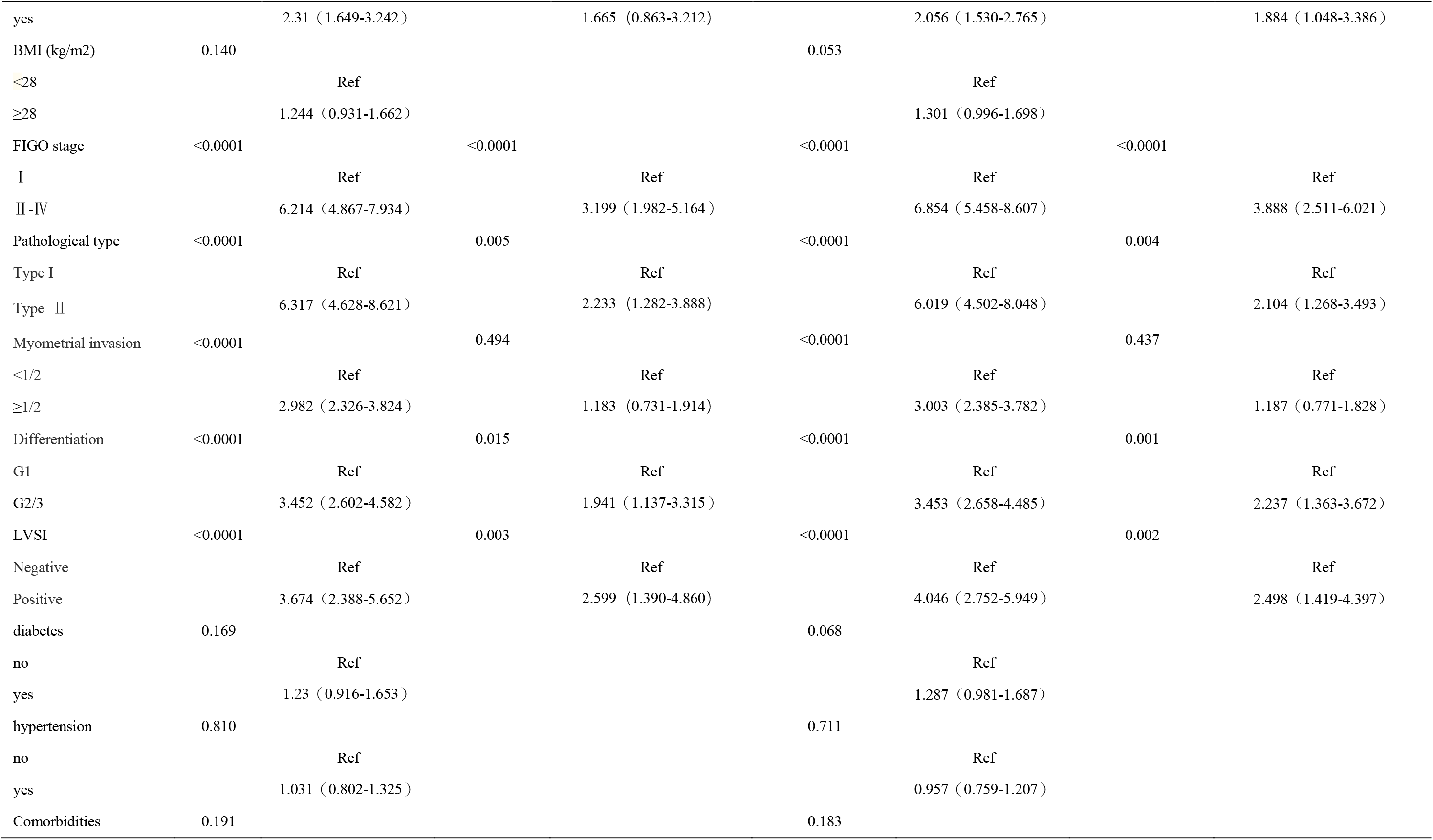
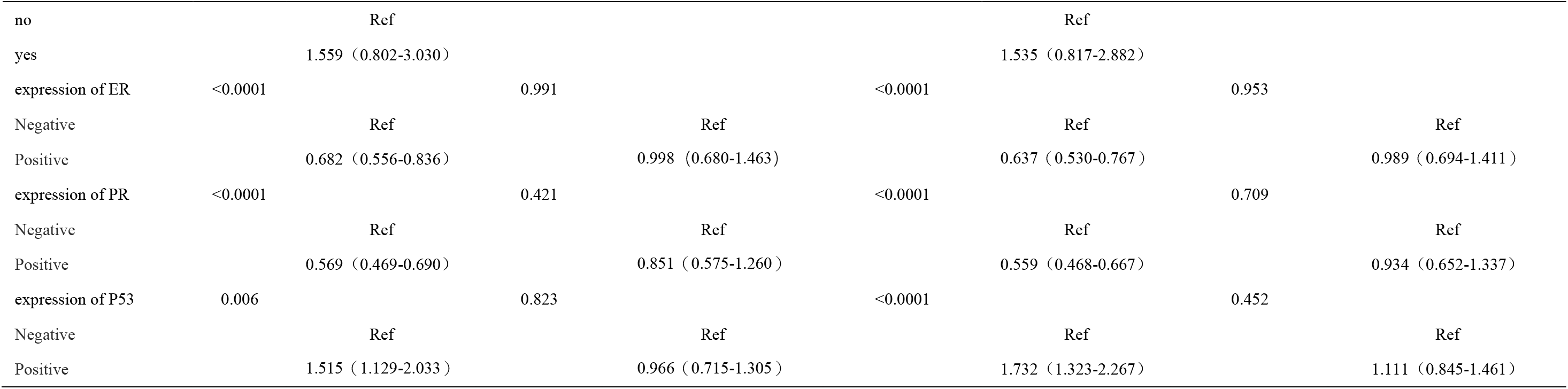
Overall survival and progression-free survival of the preoperative CA125, CA19-9, CA72-4, AFP and CEA with other clinicopathological variabl.

A multivariate analysis of factors that had been identified as being important by univariate analysis was performed (Table 2). These factors included age, menopausal status, pathological type, degree of differentiation, FIGO stage, depth of myometrial invasion, lymphovascular invasion (LVSI), expression of ER, PR, P53 in pathological tissues, serum CA125, CA72-4, CEA, and AFP levels. Elevated serum CA125 (P < 0.001) and AFP (P < 0.001), CEA (P = 0.037), age (P = 0.023), menopausal status (P = 0.034), pathological type (P = 0.004), degree of differentiation (P = 0.001), FIGO stage (P < 0.001), and lymphovascular infiltrates (P < 0.001) were independent factors for PFS prognosis in EC patients. Increased serum CA125 (P = 0.003), AFP (P < 0.001), CEA (P = 0.014), age (P = 0.004), pathological type (P = 0.005), degree of differentiation (P = 0.015), FIGO stage (P < 0.001), and lymphovascular infiltrates (P = 0.003) were independent factors for prognosis of OS in EC patients (Table 2).

### 3.4 Construction of risk score based on serum CA125, CEA, AFP level to predict patient outcomes

To further investigate the role of the platelet indexes in prognosis prediction, the CA125, CEA, AFP levels were combined to establish the risk score. EC patients included in the study were randomly divided into a test cohort and a validation cohort according to a 1:1 ratio (Table 3). In the training cohort, the least absolute shrinkage and selection operator (Lasso)-Cox regression model was used to screen the independent factors for OS in EC patients (Fig. 4). CA125, AFP and CEA were screened as the best modeling markers to construct the EC prognostic risk score model. The risk score of each patient was calculated according to the regression coefficient and serological level of the three serological tumor markers. Risk score =0.000382628046162818×level of CA125+0.00321692637528874×level of AFP+ 0.0269225927239563×level of CEA. The risk score of each patient was calculated according to the above risk scoring formula, and the ROC curve was drawn (Fig. 4). The sensitivity and specificity corresponding to different risk probabilities were calculated using the ROC curve in the training cohort, and the Yoden index corresponding to different risk probabilities was further calculated (Yoden index = sensitivity + specificity –1). The maximum Yoden index was used to determine the optimal risk threshold of OS predicted by the model, which was 0.0717. According to this, patients were divided into high and low risk groups, and the risk curves of the two groups were drawn. The death rate of patients in the high risk group was higher, indicating that the high risk group was more likely to have poor prognosis. In the verification cohort, the ROC curve of the drawing time shows that the AUC value of 1-5 years in the verification cohort ranges from 0.740 to 0.808, indicating that the above model still has good prediction performance in the external verification cohort. The risk score of each sample was calculated according to the above risk scoring formula (Fig. 4), and the Kaplan-Meier survival curve was drawn, indicating that the survival rate of the high-risk rating group was significantly lower than that of the low-risk rating group (P < 0.0001).

**Figure 4.** (A): Screening prognostic factor by Lasso regression analysis, Cross-verify the confidence interval distribution of the variable and the target variable; (B): Receiver operating characteristics (ROC) curve analysis of the Risk-score for OS in training cohorts; (C): Kaplan– Meier curves of OS for each Risk-score level group in training cohorts; (D): Receiver operating characteristics (ROC) curve analysis of the Risk-score for OS in validation cohorts; (E): Kaplan– Meier curves of OS for each Risk-score level group in validation cohorts.

**Table 3:**
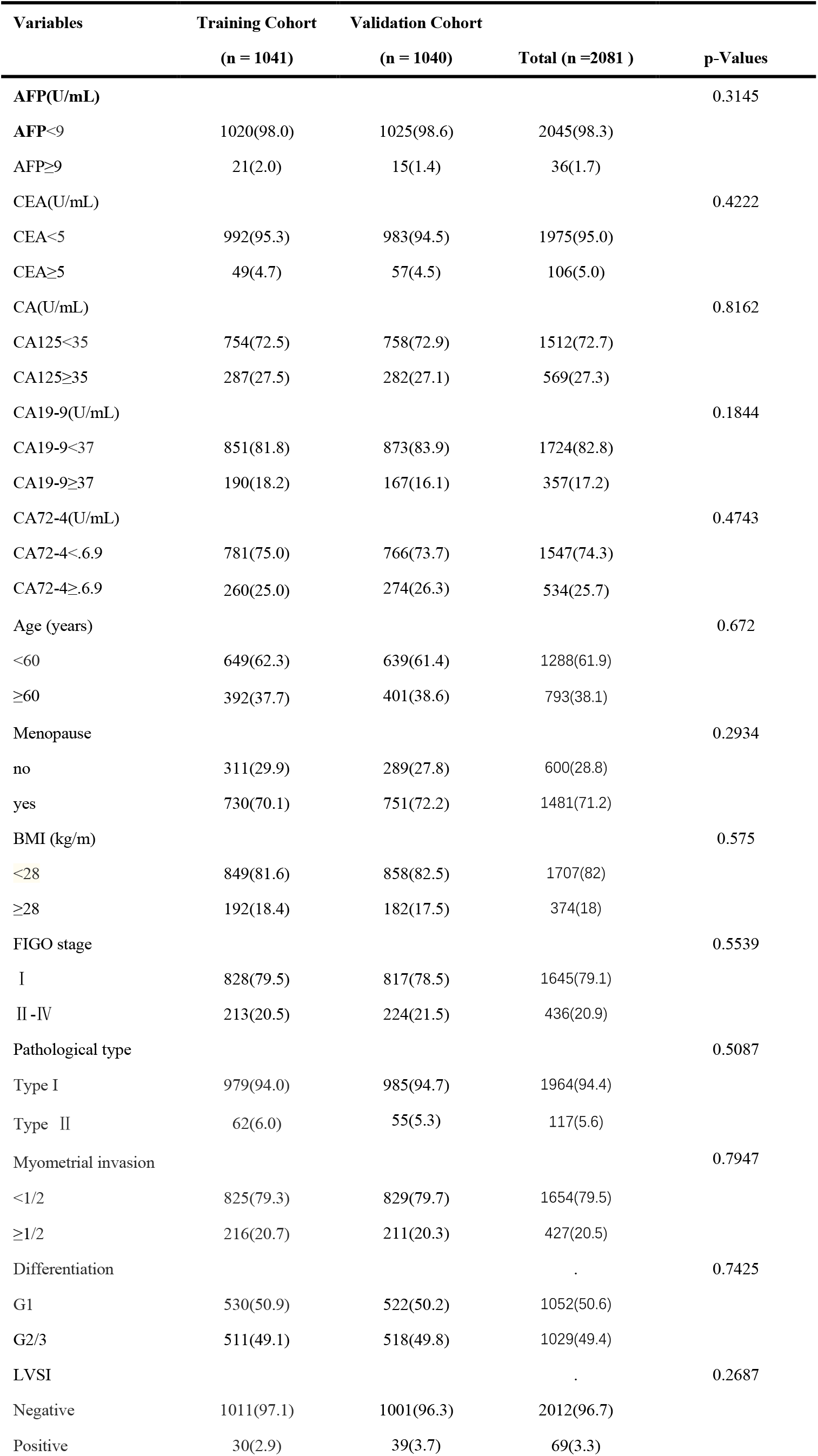
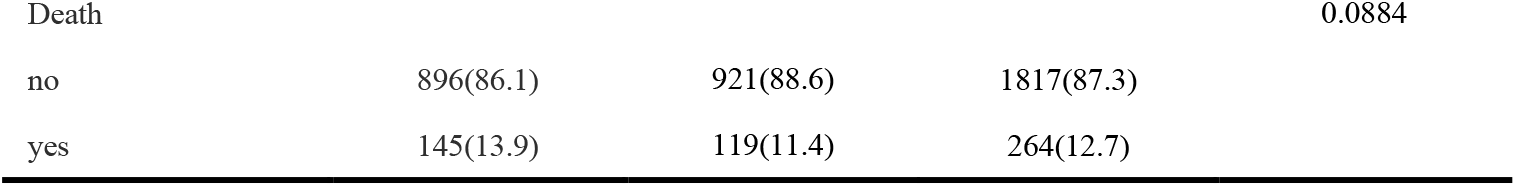
Patient biomarkers and characteristics of the training and validation cohorts.

In the entire EC cohort, risk score, age, grade, FIGO stage, pathological classification and other clinical information were integrated into the Univariate and Multivariate Cox regression analysis of OS and PFS (Table 4). Univariate analysis showed that there was a statistical difference in risk scores between OS and PFS (P<0.0001). Kaplan-Meier survival curve showed that, the survival time of PFS and OS in high-risk rating group was significantly lower than that in low-risk rating group (p <0.0001) (Fig. 5). Combined with Multivariate Cox regression analysis, RS was an independent prognostic factor for patients with PFS and OS (P<0.0001) and could be independently used as a prognostic indicator for patients with EC (Table 4).

**Figure 5.** (A): Kaplan–Meier curves of OS for each Risk-score level group in all endometrial cancer patients; (B): Kaplan–Meier curves of PFS for each Risk-score level group in all endometrial cancer patients.

**Table 4:**
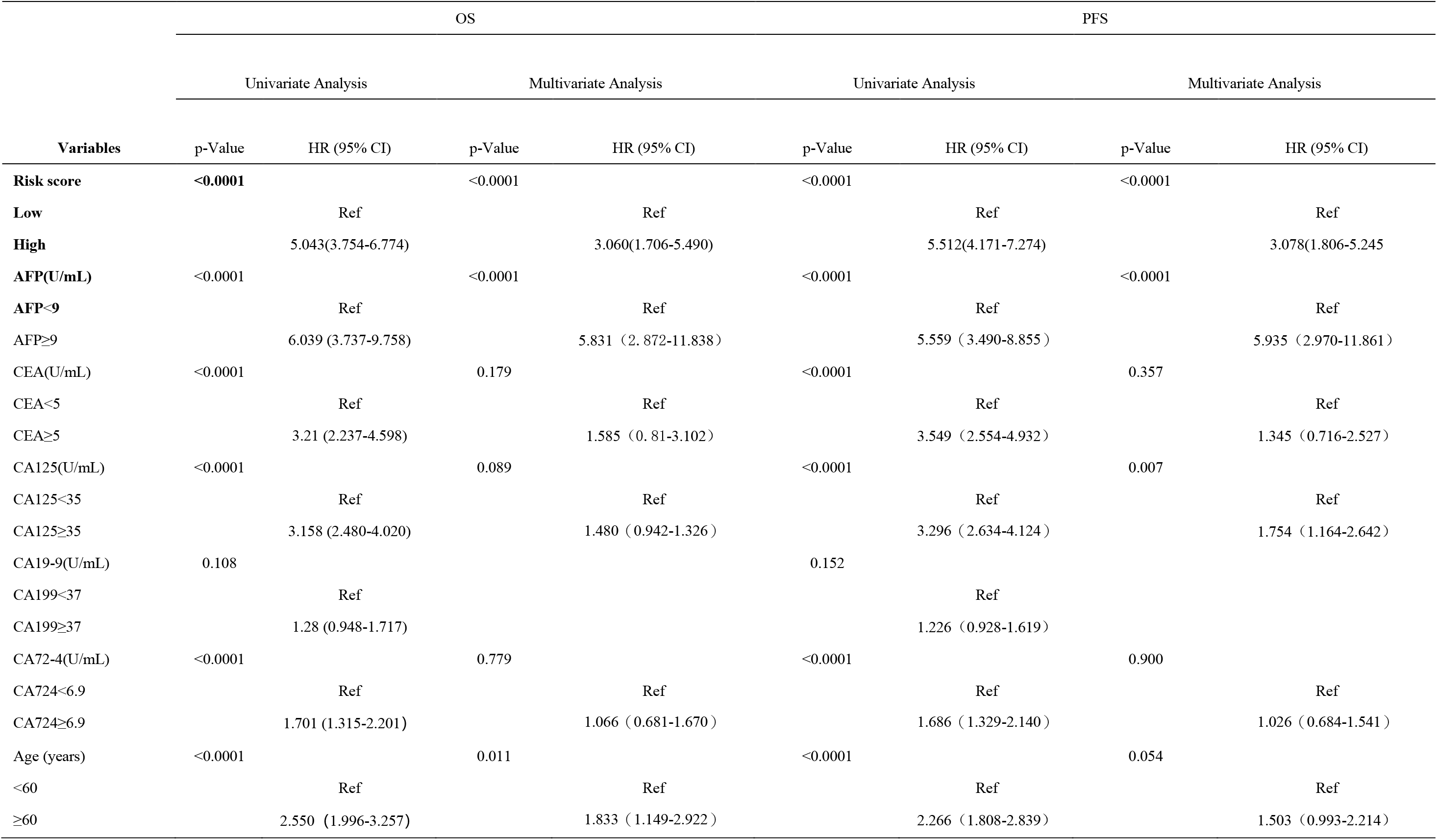
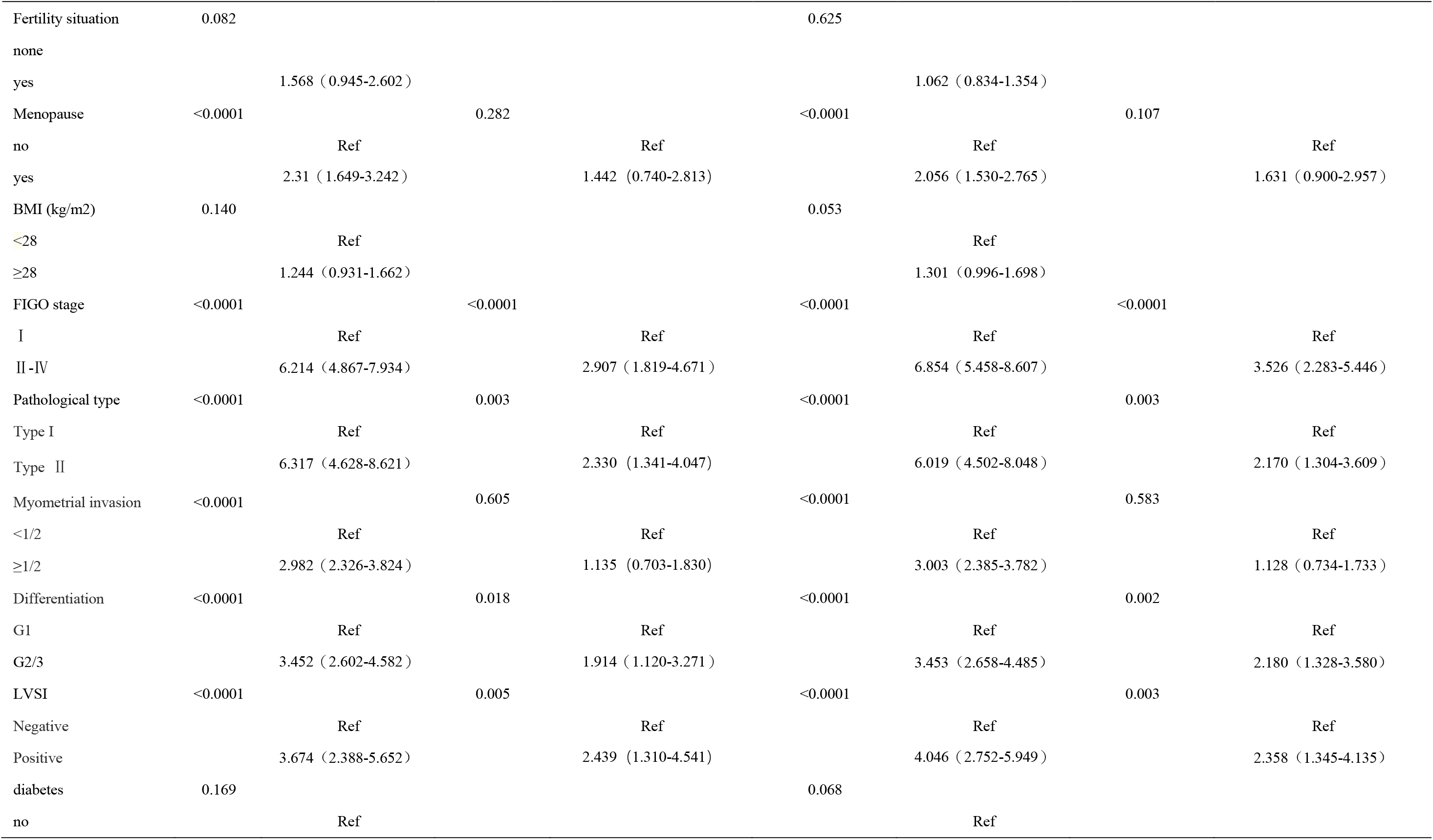
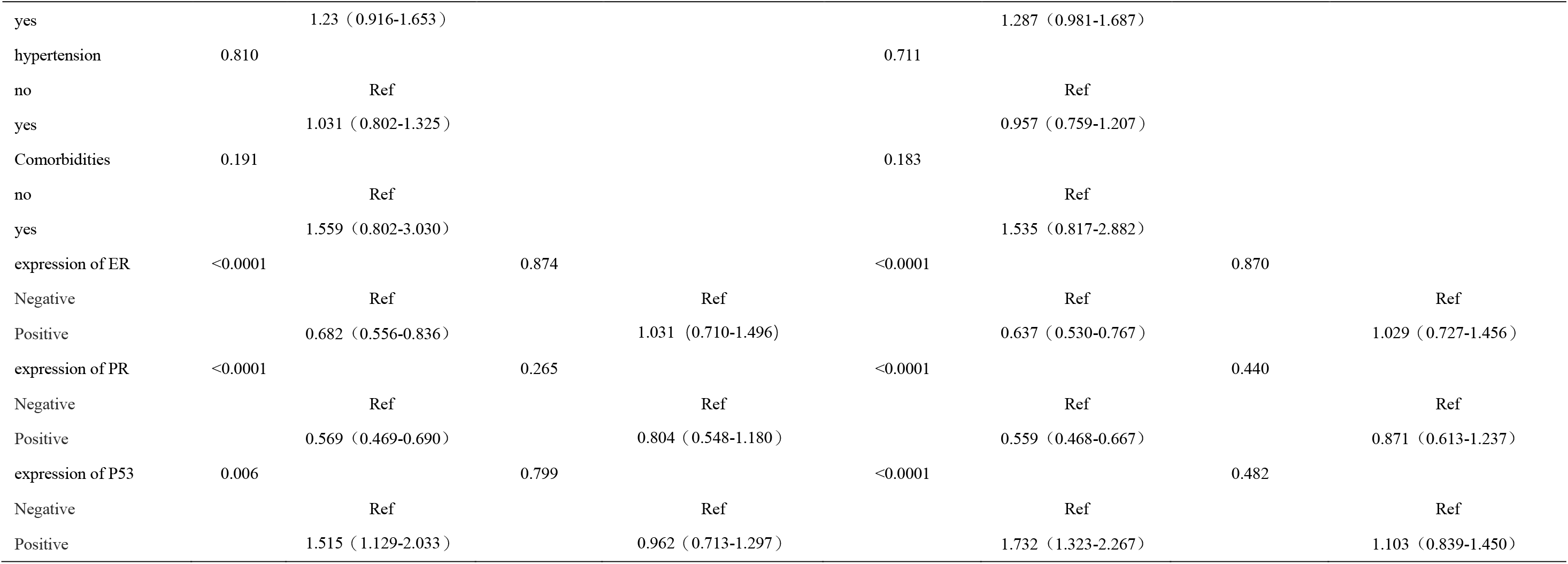
Overall survival and progression-free survival of the Risk score with other preoperative biomarkers and clinicopathological variables.

### 3.5 Nomogram Construction and Validation

Based on the results of the Cox regression models in Table 2, nomograms were constructed by integrating the independent prognostic factors for PFS and OS (Fig. 6). In addition to the risk score, FIGO stage, Age and degree of differentiation was included in the nomograms, Summing the scale points corresponding to each variable on the nomogram corresponds to the total score on the bottom scale, which represents the 3– and 5-year PFS and OS probabilities that were predicted by the model. the C-index of the multivariate prognostic model of PFS according to the risk score, age, FIGO stage, and differentiation variables was 0.836 (95% CI = 0.822–0.918). The C-index of the multivariate prognostic model of OS according to the risk score, age, FIGO stage, and differentiation variables was 0.837. The C index showed that the prediction results of the prognostic model had a high probability of agreement with the actual observation results. These findings indicated that the model had good discriminative ability to separate patients with different survival outcomes. The calibration plots of the nomogram show good consistency between the actual observations and the predicted probabilities of PFS and OS (Fig.7).

**Figure 6.** Nomograms for predicting the postoperative PFS (A) and OS (B) for 1, 3 and 5 years in endometrial patients.

**Figure 7.** (A): Calibration plots of the nomograms of PFS for 1 year in endometrial cancer patients;(B): Calibration plots of the nomograms of PFS for 3 year in EC patients;(C): Calibration plots of the nomograms of PFS for 5 year in EC patients;(D): Calibration plots of the nomograms of OS for 1 year in EC patients;(E): Calibration plots of the nomograms of OS for 3 year in EC patients;(F): Calibration plots of the nomograms of OS for 5 year in EC patients;

## 4. Discussion

EC is one of the most common female malignancies as well as the third most universal female genital malignancy. While most patients are diagnosed relatively early, a significant number of patients are still being diagnosed in the later stages of disease. The FIGO staging system has guiding significance for the assessment of prognosis and treatment of EC patients; however, prognostic status could vary significantly even for patients divided into the same stage.

In our study, We included 2081 patients with endometrial cancer undergoing surgical treatment to perform an analysis of independent prognostic factors. Patient groups with high and low levels of CA125, CA199, AFP, CEA and CA724 were established, High level of CA125 is considered as a risk factor for poor prognosis of endometrial cancer patients, which is consistent with previous studies. It is worth mentioning that in our patient cohort, AFP and CEA were also identified as independent prognostic factors for endometrial cancer patients, and patients with high levels of AFP and CEA showed poor prognosis. Therefore, we constructed a new endometrial cancer risk parameter using AFP, CEA and CA125 levels to predict the clinical prognosis of PFS and OS in endometrial cancer patients.

Least absolute shrinkage and selection operator (LASSO) is a regression analysis method, which can simultaneously select and regularize variables to improve the prediction accuracy and interpretability of the generated statistical model. The algorithm has been widely applied to Cox proportional risk regression model (23, 24), and is used for survival analysis of high-dimensional data. Lasso Cox regression can be used to screen out independent variable sets with strong explanatory power for dependent variables, which is helpful to improve the refinement and accuracy of the prediction model. It is mainly applied to the selection of variables with small samples and multiple factors. This study used LASSO regression model to explore the prognostic factors of endometrial cancer patients. Based on multiple serum tumor markers, this risk score showed high efficacy in predicting patient outcomes, and patients in the high-risk group assigned by this score showed worse PFS(HR=3.078 95% CI=1.806-5.245) and OS (HR=3.060 95% CI=1.706-5.490). The risk profile of this score may be a more targeted and powerful prognostic assessment for predicting positive clinical outcomes, and may be a more effective classification tool for EC patients. We further constructed a nomogram using risk score, FIGO staging, grading, and patient age. The c index of the model is greater than 0.8, and the calibration curve is close to the diagonal, indicating that the model has good predictive ability to determine the prognostic value of this combination for EC patients.

Tumor biomarkers are substances expressed and secreted in the process of tumorigenesis and development. They may indicate the existence of new tumor growth, which means relatively large tumor burden and invasive phenotype. Serum tumor biomarkers are widely used in clinic and provide important information for early diagnosis, evaluation of curative effect, prognosis and prediction of metastasis and recurrence of malignant tumors. The measurement of serum tumor biomarkers is convenient and economical. Therefore, they are promising tools for guiding treatment plans and long-term monitoring of recurrence and metastasis. The area of research investigating biomarkers for EC is relatively well established(*25–27*). Niloff et al. first reported that serum CA125 was elevated in women with recurrent and advanced endometrial cancer in 1984(*13*). Subsequent studies have consistently shown that there is a correlation between serum CA125 concentration and the clinicopathological parameters and outcomes of poor endometrial cancer(*28–31*). CA125 can also be used as a preoperative indicator to evaluate whether regional lymph nodes are invaded, which can predict OS and DFS. In these studies, CA125 is mainly used to monitor the postoperative recovery and long-term prognosis of EC patients(*29, 32, 33*), or to improve the EC positive rate in combination with HE4. In this study, both univariate and multivariate analyses showed that elevated serum levels of CA125 indicated a poor prognosis for OS and PFS in EC patients. Patients with serum concentration of CA125 ≥ 35.0 mM had a 1.9-fold higher risk of death due to cancer compared to patients with baseline levels of CA125 (P = 0.003, HR =1.989, 95% CI: 1.270-3.116). This is consistent with previously published literature CEA is an acidic glycoprotein with the characteristics of human embryonic antigen. It exists on the surface of cancer cells differentiated from endodermal cells and belongs to the structural protein of cell membrane. It is synthesized in the small intestine, liver and pancreas during the embryonic stage, and the content of CEA in adult serum is extremely low. CEA was first used as a specific marker for the diagnosis of colorectal cancer(*34*). The dynamic detection of serum CEA has important reference value for the judgment of recurrence or metastasis of gastrointestinal cancer(*35*), and CEA can also be used as a good indicator for the prognosis evaluation of colorectal cancer. In recent years, research shows that CEA has certain reference value in the diagnosis of female reproductive system tumors(*36*). A study by Gadducci A et al. shows that CEA is also overexpressed in elderly endometrial cancer patients(*37*). Although it cannot be used as a specific indicator of elderly endometrial cancer, it plays an important role in the diagnosis of differentiation of endometrial tumor and evaluation of curative effects. This study is the first to describe the potential use of serum CEA as a prognostic tool for EC. Patients with elevated serum CEA are more likely to have worse outcomes than patients with normal concentration of serum CEA.

AFP is a glycoprotein that is mainly synthesized by fetal liver cells and the yolk sac. It is found in high concentrations in the fetal circulatory system and declines shortly after birth to be functionally replaced by albumin. It is present in extremely low concentrations in adult serum(*38*). AFP has many important physiological functions, including transport functions, bidirectional regulation of growth, immunosuppression, and is also involved in T cell-induced apoptosis. AFP is the most widely used serological marker for the diagnosis of hepatocellular carcinoma(*39*). It is also elevated in a variety of extrahepatic tumors, including yolk sac tumors, gastrointestinal tumors, and pancreatic, gallbladder, lung, and bladder cancers(*40–42*). This study is the first to describe that serum AFP can be used as an independent risk factor for the prognosis of EC. AFP measurement may provide further information after surgical risk stratification to help clinicians predict prognosis and provide basis for clinical treatment. Studies have found that endometrial carcinoma producing AFP is a clinicopathologic subtype of endometrial carcinoma(*43–45*), which occurs in 1.8% of endometrial carcinoma cases(*46*). This classification is related to TP53 abnormality and vascular invasion. Even in FIGOI cases, these cancers can show aggressive behavior(*45*). Similarly, in our study cohort, 1.73% of patients showed AFP expression above the normal range, and this part of patients also showed significant adverse prognosis. we identified serum concentration of AFP ≥9ng/ml in EC patients as a marker for poor prognosis for PFS (P<0.0001, HR =7.437, 95% CI: 3.717-14.879) as well as OS (P< 0.0001, HR = 6.914, 95% CI: 3.374-14.165).

The study still has limitations. Some patient accurate information was unable to obtain due to their loss of follow-up. However, as a large EC patients sample population was enrolled, and the data from the electronic medical record system of Shengjing Hospital Affiliated to China Medical University were verified, this provides a firm basis for the conclusions drawn. In conclusion, AFP concentration is a promising biomarker for EC and a valuable independent factor for predicting patient prognosis and survival. levels of serum CA125, AFP, and CEA concentration were promising biomarkers for EC and a valuable independent factor for predicting patient prognosis and survival. However, a risk score created by combining CA125, AFP, and CEA levels to predict the prognosis of endometrial cancer was found to be highly accurate and valuable for clinical use.

## Author contributions

Zi-hao Wang participated in the design of the work, methodology, data interpretation, and analysis for the work; and drafted the manuscript. Yun-zheng Zhang and Bo Wang Wang carried out the statistical analyses and drafted the manuscript. Shu-wen Ge, Lu-he Shan and Zi-yu Zhang participated in the methodology, data interpretation, and analysis for the work. Qi-jun Wu and Xiao-xin Ma designed the study; participated in data interpretation, analysis for the work, and methodology. All authors read and approved the final manuscript.

## Abbreviations

EC: endometrial cancer
OS: overall survival
PFS: progression-free survival
FIGO: International Federation of Gynecology and Obstetrics
CA125: carbohydrate antigen 125
CA19-9: carbohydrate antigen 19-9
CA72-4: carbohydrate antigen 72-4
AFP: alpha-fetoprotein
CEA: carcinoembryonic antigen
HE4: human epididymal protein 4
HR: hazard ratio
CI: confidence interval

## Acknowledgements

This work was supported by the National Natural Science Foundation of China (No. 81872123 and 81472438); University innovation team of Liaoning Province; Special Professor of Liaoning Province; “Major Special Construction Plan” for Discipline Construction of China Medical University in 2018(No. 3110118029); Outstanding Scientific Fund of Shengjing Hospital(No. 201601) “Major Special Construction Plan” for Discipline Construction of China Medical University in 2018.

## Availability of data and materials

There are no linked research data sets for this paper. Data are available from the corresponding author on reasonable request.

## Consent for publication

All listed authors have actively participated in the study and have read and approved the submitted manuscript.

## Ethics approval and consent to participate

This study was approved by the Ethics Committee of Shengjing Hospital of China Medical University.

## Conflicts of Interest

The authors declare no conflict of interest.

## Notes

### Competing Interest Statement

The authors have declared no competing interest.

